# Streamlining large-scale high-resolution electron tomography with VolWeaver

**DOI:** 10.64898/2026.08.14.744809

**Authors:** Irina Bregy, Rob Mesman, Silvia Tassan-Lugrezin, Taco W. A. Kooij, Laura van Niftrik

**Affiliations:** Department of Microbiology, Faculty of Science, Radboud University, 6525AJ Nijmegen, The Netherlands; Department of Medical Microbiology, Radboud University Medical Centre, 6525 GA Nijmegen, The Netherlands

**Keywords:** electron tomography, volume EM, image processing, TEM, *Plasmodium falciparum*, mitochondria

## Abstract

Researchers using electron microscopy must often balance a trade-off between obtaining high-resolution structural information and preserving sufficient cellular context. At one end of this spectrum, single particle cryo-electron microscopy and cryo-electron tomography provide near-molecular detail but are typically limited to relatively small fields of view. At the other, volume electron microscopy approaches, such as scanning electron microscopy of resin-embedded specimens, capture large cellular volumes but generally at lower resolution. Consequently, linking nanoscale structural information to larger cellular architecture remains a significant challenge. To address this gap, we optimised a transmission electron tomography workflow for resin-embedded malaria parasites that allows us to visualise targeted regions of interest at nanometre-scale resolution while retaining several micrometres of surrounding cellular context. Here, we present our current best-practice pipeline for sample preparation, tomogram acquisition, and reconstruction. In addition, we introduce VolWeaver, a data-processing framework, that integrates high-resolution tomographic datasets into serial section volume reconstructions, enabling the visualisation and interpretation of ultrastructural features within their broader cellular environment.

## INTRODUCTION

### Imaging modes

With a long-standing tradition in capturing the fine details of biological ultrastructure, transmission electron microscopy (TEM) is famous for its focus on the very small. Nevertheless, the research community is well aware of the relevance of cellular context, hence often hesitant to decide between maximising local resolution and preserving information on cellular context (*1*, *2*). Resolution is typically higher in cryogenic workflows where the specimen itself (rather than a stain accumulating around it) provides contrast. Single particle cryo-TEM (SPA) and cryo-TEM tomography have experienced growing attention throughout recent years and both resolution as well as speed and ease-of-use have profited immensely from the growing interest in these methods (*3–10*). While SPA depends on repetition; imaging many, near-identical objects and then combining them computationally (*11*, *12*), cryo-TEM tomography is based on the acquisition of a tilt-series at regions of interest (ROIs), followed by computational reconstruction of 3D volumes by back projection (*13–16*). Repetition of substructures within or between the resulting 3D volumes are often used to increase local resolution similarly to the computational workflow used for SPA data processing. This additional processing step is known as subtomogram averaging. For large, abundant protein complexes subtomogram averaging can increase local resolution to near-SPA levels (*16–18*).

In contrast, high volumes are usually imaged in scanning electron microscopy (SEM) rather than TEM, a field that is currently dominated by focused ion beam (FIB) SEM, but also features mechanical shaving of resin-embedded samples using serial block face (SBF) SEM (*19*, *20*). In SEM applications, the specimen surface is scanned, shaved off and scanned again, ultimately rendering a series of surface scans that cover the full depth of the ROI (*21*). Because SEM resolution is limited by the size of the scanning probe and the thickness of the layers that are shaved off between scans, throughput and the achieved resolution are directly opposing each other (*20*, *22*).

Lastly there is the hybrid approach called scanning transmission electron microscopy (STEM) tomography, which combines the scanning approach of the SEM with tilt-series acquisition and computational back projection (*23–25*). Compared to TEM tomography, STEM tomography allows for much thicker samples of up to 1µm (as compared to 200-300nm for TEM tomography) (*23*, *26*). However, just like in SEM, STEM resolution depends on the size of the electron probe.

### Sample preparations

In recent years, many of the advancements in the electron microscopy field have been focusing on the refinement and expansion of the toolset available for cryogenic applications. Important advantages of working in cryogenic conditions on vitrified samples is that specimen manipulation is minimal, usually restricted to a single, rapid freezing step (*27*). In vitreous ice, sample contrast solely relies on the phase contrast provided by the specimen itself, thus avoiding artefacts of staining, dehydration and resin embedding (*28*). However, a major downside of vitrified specimen inside an electron microscope is that these samples are very fragile to an extent that prolonged exposure to the electron beam rapidly degrades the sample by local evaporation of the melting ice (*29*). As a result of this fragility, electron dose must remain low, reducing the achievable contrast. Especially tomographic acquisitions suffer from this trade off, as the total tolerable electron dose must be spread over dozens of micrographs that are acquired per ROI. Furthermore, it is worth noting that re-imaging of a pre-exposed area is not possible, as the area will already be damaged. The operator depends on the lucky shot with limited insights into the quality and biological properties of the chosen ROI and overlapping montages of neighbouring regions are not feasible.

Sample instability is largely a result of the presence of water in the sample. While frozen, the water remains in place despite the vacuum (*28*). Local interaction with the electron beam, however, causes the water to evaporate, hence causing bubble formation (*29*). For this reason, truly stable sample preparations that withstand the electron beam are free of water and thus require dehydration, staining and resin embedding (*30*). To remove water from the sample, researchers use stepwise replacement with organic solvents such as ethanol or acetone (*30*, *31*). During or after this process, the sample has to be stained with heavy metals to increase amplitude contrast (*30*). The sample is then infiltrated with resin to generate a block of mechanically stable material (*31*, *32*). Such resin-embedded specimen can be used for SBF-SEM or FIB-SEM directly or cut in serial or individual sections. The added amplitude contrast of these preparations is a disadvantage as much as it is an advantage. While it greatly increases visibility of structures of interest, it comes at a resolution cost and potentially introduces artefacts (*33*). Such artefacts may be introduced during sample fixation, dehydration, staining and sectioning. To prevent fixation and dehydration artefacts, scientists often resort to high-pressure freezing, a method that is used also in cryogenic preparations. It usually allows for skipping chemical fixation and more importantly, it allows for more gentle removal of water during the downstream process of freeze-substitution (*34–36*).

### Large volume acquisitions

High-resolution and large-scale acquisitions often come at a trade-off, where resolution decreases with larger fields of view (*37*). Most commonly, FIB-SEM is used to achieve rather high resolution of large areas (*20*, *22*). However, most studies using FIB-SEM do not exceed resolutions of ∼5nm due to probe size limitation and FIB-laser increment. Highest resolutions are typically achieved at high magnification in TEM, especially cryo-electron tomography (cryo-ET) (*38*). Recent studies introduced the concept of serial lift out cryo-FIB milling to recover large z-stacks for cryo-ET (*39*).

Despite the aforementioned cost in resolution and risks of artefact introduction however, it is quite common to use resin-embedded samples to gather large-scale volumetric data (*40*, *41*). One substantial advantage of resin-embedded samples is their stability in the electron beam. This offers the possibility for multiple acquisitions at the same ROI, either on overlapping subregions or at different magnifications. Our recent work on *Plasmodium falciparum* (the etiological agent of the most severe form of human malaria) demonstrates that the resolution achieved with optimised sample preparation and acquisition setup is sufficient for tracing membrane architecture throughout the cell (*42*). Here, we present the workflow that allows us to supplement low-resolution overview tomograms with high-resolution insets. Serial sectioning of our chemically prefixed, high-pressure frozen, freeze-substituted and resin-embedded samples allows us to collect and integrate targeted high-resolution information across depths of several microns. For our research, we generated a python-based interactive software (VolWeaver) that features semi-automatic montaging and embedding of high-resolution volumes into a larger canvas of low-resolution cellular context and serial joining of z-adjacent volumes. Furthermore, VolWeaver comes with a set of modelling tools to aid manual segmentation or refinement of automatic segmentations performed by any third-party software. The application also offers a range of conversions between common data formats in the electron microscopy field, filtering options as well as a set of measurement tools that we designed for our research on mitochondrial cristae.

## RESULTS AND DISCUSSION

### Combining cryogenic fixation with chemical fixation to prevent stress-related changes in *Plasmodium falciparum* blood-stage parasites

Malaria parasites are prone to temperature-related morphological changes, exemplified by male gametocytes, which rapidly initiate exflagellation following a drop in temperature and increase in pH upon transmission to the mosquito midgut (*43*). Biosafety concerns further complicate parasite fixation by high-pressure freezing. High-pressure freezing of freshly harvested *P. falciparum* must be performed in the BSL2 laboratory following established safety protocols (*44*).

In the setup available to us, ensuring biosafety while avoiding parasite stress was challenging to the point that, despite meticulous planning, we observed severe morphological damage and signs of mitochondrial stress in *P. falciparum* gametocyte samples frozen directly from culture (Figure 1A). In contrast to the well-documented elongated banana shape of mature gametocytes (*45*), the parasites appeared deformed, containing ruffled membranes and distorted organelles. Mitochondria frequently contained what appeared as holes or double-membrane invaginations and multi-membrane structures of unknown identity were observed. Thus, we decided to chemically prefix our samples prior to high-pressure freezing and subsequent freeze-substitution. This protocol was chosen based on the rationale that dehydration at room temperature may cause shrinkage and thus affect membrane architecture (*46*, *47*). The fact that cellular membranes appeared smooth in the generated TEM samples implied that our protocol preserved native membrane architecture (Figure 1A). Furthermore, while cell and organelle shape appeared distorted in samples frozen directly from culture, the morphology observed in chemically prefixed parasites matched the expected elongated banana-shape of mature *P. falciparum* gametocytes (*45*, *48–50*). We compared two different chemical fixation cocktails (4% paraformaldehyde (PFA) in PHEM buffer versus 1% PFA + 0.0075% glutaraldehyde (GA) in MOPS-buffered saline (MBS)) and did not observe any differences in cell preservation (compare Figure 1A with Figure S 1A). Though equally suited for pure electron microscopy studies, we mainly used the fixation mix consisting of 1% PFA + 0.0075% GA in MBS for our later experiments.

**Figure 1:**
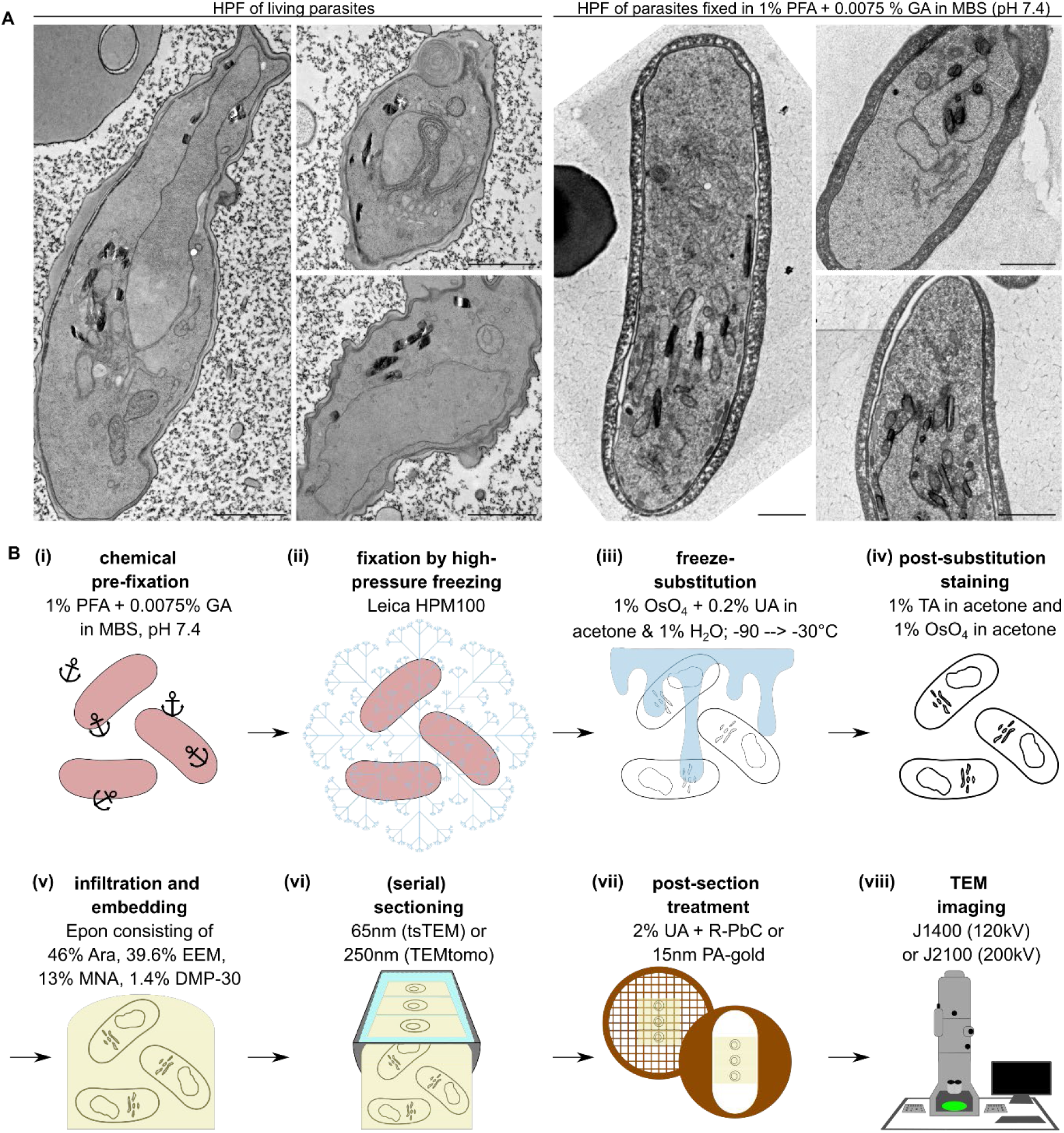
Plasmodium falciparum sample preparation workflow – from the living parasite to the microscope. A) Comparison of P. falciparum gametocytes when high-pressure frozen directly from culture, versus prefixed with PFA and GA and then high-pressure frozen. Both samples were subsequently freeze-substituted and resin-embedded. Scale bar 1µm, TEM of 65nm thin sections. B) Overview of sample preparation workflow for TEM. Our current workflow starts with a chemical prefixation (i) followed by high-pressure freezing (ii), and freeze-substitution in OsO_4_ and UA dissolved in acetone with 1% water (iii). Post-substitution staining consists of TA treatment followed by another layer of OsO_4_ (iv). Samples are then infiltrated with an Epon mix optimised to match intrinsic sample hardness (v). Hardened Epon blocks are sectioned (serially) at a diamond knife (vi). For recovery of whole cells by TEM tomography (TEMtomo), 10-15 250nm sections are collected on a single-slot grid (vii). For thin section TEM (tsTEM), 1-5 65nm sections are collected on meshed grids. Either section-type was imaged at one of our in-house TEMs (viii). For sufficient contrast on high tilts of 250nm sections, we recommend a 200kV microscope.

Guided by pre-existing literature on sample preparation strategies, we tested the inclusion of small amounts of water in the freeze-substitution mix (*51*) and downstream membrane enhancement with tannic acid (TA) treatment (*52*). Our results point towards a sweet spot at 1% ultrapure water in a mix of acetone in combination with standard concentrations of heavy metals and post-substitution staining with TA (Figure S 1B). Temperature slopes and incubation times were not further optimised in the scope of this study.

Our gold standard for sample treatment thus consisted of (i) chemical prefixation, (ii) high-pressure freezing, (iii) freeze-substitution alongside osmium tetroxide (OsO_4_) and uranyl acetate (UA) staining and 1% ultrapure water for optimal membrane contrast, (iv) post-substitution staining with TA and additional OsO_4_ impregnation, (v) infiltration and embedding in epoxy resin (Epon), (vi) serial sectioning, (vii) post-section treatment and (viii) imaging at the TEM (Figure 1B).

### Data acquisition at high resolution with large cellular context window

To capture cellular context across multi-micron depths, we collected serial sections at 250nm thickness and collected them on single-slot carbon-formvar coated copper grids (sample processing overview in Figure 3A). Per grid we attempted to collect as many sections as possible, typically between 10 and 15. We then recorded low-magnification overview tilt-series across the entire cell, following the ROI through the adjacent sections. To speed up this process, we collected the low-magnification tilt-series at our screening microscope at 120kV, using relatively large increments (2-3°), and limited tilt-range to ± 45°, but retaining the dual-axis routine. From the reconstructed overview tomograms, we could clearly identify the areas where more resolution was desirable, and selected these ROIs for imaging at higher magnification, tighter increment, wider tilt-range and at 200kV. We automated acquisition using SerialEM and reconstructed all tomograms with the IMOD batch reconstruction workflow (*53*, *54*). This allowed us to acquire and reconstruct large datasets with minimal active work time. Examples of our tomographic data are shown in Figure 2 as well as in our recent work on the mitochondrial contact site and cristae organising system in *P. falciparum* (*42*).

**Figure 2:**
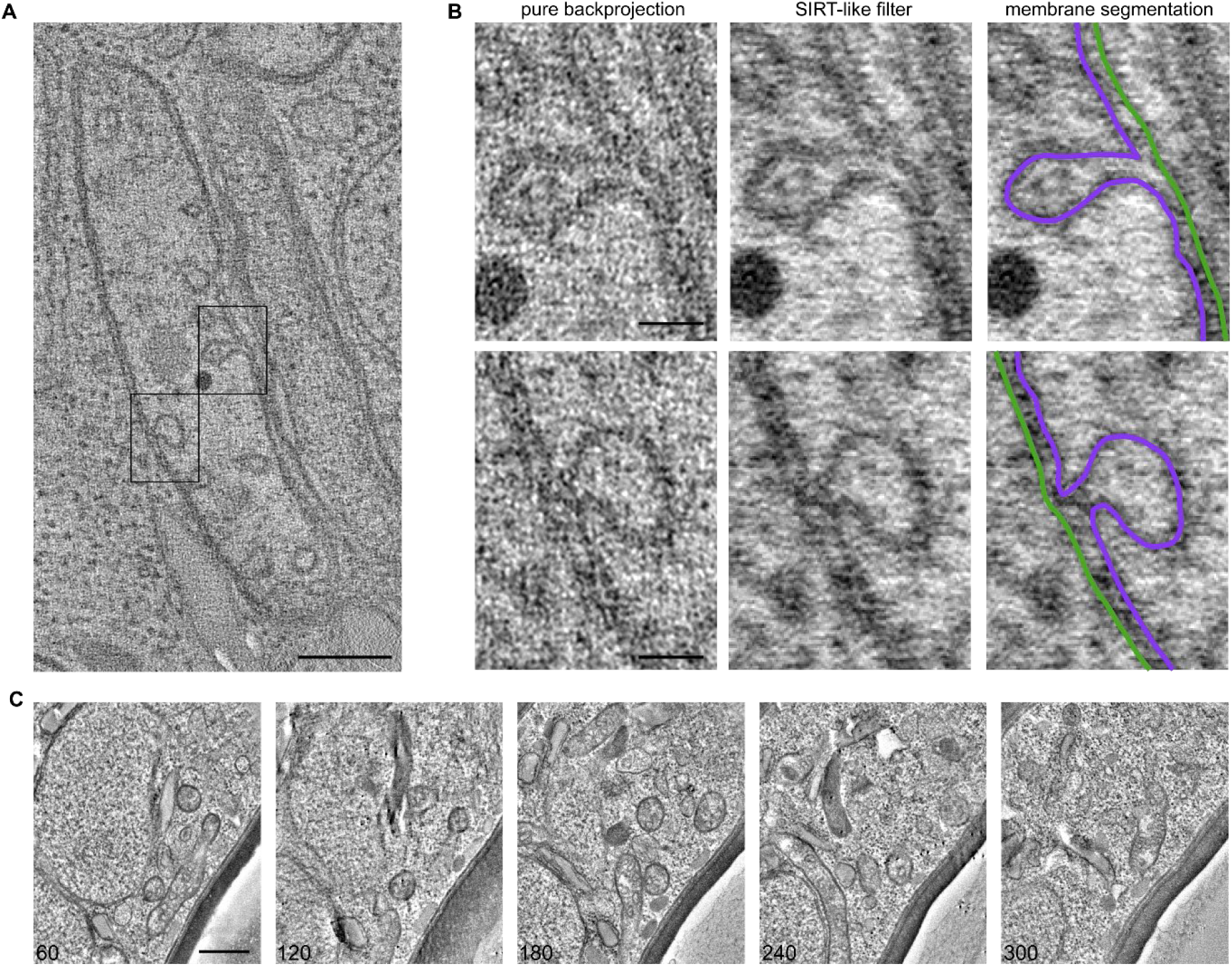
Snapshots from tomographic volumes of Plasmodium falciparum gametocyte mitochondria. A) snapshot derived from a tomogram acquired on a 250nm section of prefixed, high-pressure frozen, freeze-substituted and resin-embedded P. falciparum gametocytes, with two rectangles highlighting individual cristae that are also shown in B. Scale bar: 250nm. B) zoom in of the cristae marked in A. (from left to right) pure backprojection; SIRT-like filtered (20 iterations); membrane segmentation of the inner (purple) and outer mitochondrial membrane (green). Scale bars: 50nm. C) Series of snapshots along a serial tomographic acquisition across 5 consecutive 250nm slices from prefixed, high-pressure frozen, freeze-substituted and resin-embedded P. falciparum gametocytes. Numbers indicate slice number along the combined stack. Full movie available as Movie 1. Scale bar: 500nm.

**Figure 3:**
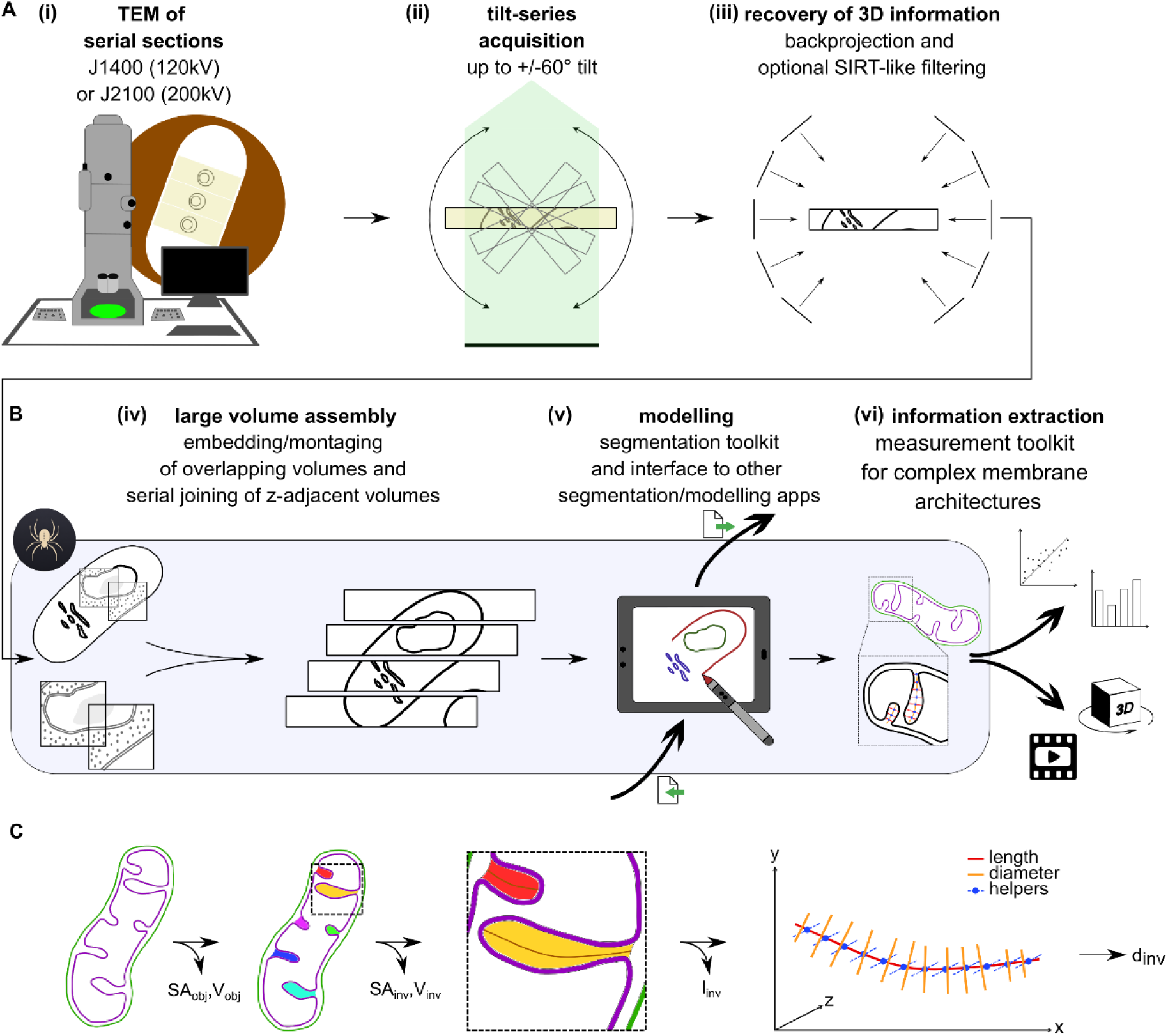
Image processing workflow – from individual tomograms to supervolumes, models, and numbers. A) Serially sectioned samples collected on single-slot grids were imaged at our in-house TEMs (i) by acquiring dual axis tilt-series of tilt-ranges up to ± 60°(ii). 3D information was recovered by backprojection and in some cases coupled with SIRT-like filtering (iii). B) We assembled low-resolution overview tomograms with high-resolution insets in VolWeaver and combined these composites to supervolumes from adjacent sections (iv). VolWeaver offers a set of segmentation tools that includes interpolations and cleanup options (v). In the VolWeaver measurements toolkit we include options to obtain volume, surface area, length, and diameter of segmented objects and substructures thereof (vi). Specifically, the program is designed to detect and measure tubular membrane invaginations automatically. C) Automatic detection and measurement of membrane invagination dimensions in VolWeaver, consisting of automatic invagination detection, automatic length path determination in tubular invaginations and incremented measurements of the diameter along each invagination. Surface area and volume per object (SA_obj_ and V_obj_) are determined from the flood-filled object outlines. Invaginations are found based on bay and hole detection and matched by z-overlap. From the flood-filled invagination mask, surface area and volume (SA_inv_ and V_inv_) are determined automatically. The diameter is determined as follows: measurement positions are placed along the invagination’s length path and z-aligned helper lines are created orthogonally to the local tangent of the length path. Helper lines are then used to guide an array of diameter measurements at each measurement position. These measurements are marching through z, measuring the xy-aligned diameter along at each helper line. The widest reasonable diameter detected at a given helper line is defined as the local diameter.

As a measure of local resolution within higher magnification acquisitions, we specifically inspected cellular membranes, which we frequently observed as sets of two parallel lines (Figure 2B). This observation serves as a standard benchmark in the field, confirming that local resolution is adequate to distinguish the two leaflets of the bilayer membrane separated by the typical 3–5 nm distance between phospholipid head groups (*55*, *56*).

### Data processing with VolWeaver

For biological interpretation, we needed to integrate information of adjacent and overlapping volumes together (example of a serial tomogram assembly in Figure 2C and Movie 1). Although workflows for serial volume assembly are available in IMOD (*53*), the management of increasingly large intermediate datasets and the absence of several workflow-specific functionalities prompted us to develop a complementary processing framework.

Thus, we decided to optimise our data processing workflow by creating VolWeaver, a data processing pipeline. With a suitable toolset for high-resolution serial and overlapping tomography, VolWeaver is designed specifically for large-scale volume electron microscopy. The python-based program is available open-source, on Github (https://github.com/irinabregy/VolWeaver). It offers workflows for all processing steps required downstream of the reconstructed tomogram, also featuring a measurement tab that allows for determining the length, diameter, volume and surface area of substructures within a membrane segmentation map, as well as per object surface area and volume determination. Importantly, VolWeaver integrates output files from other sources, including automatic segmentations done with third-party tools and applications. It accepts both MRC masks and IMOD-style models, converts between them and offers manual drawing tools as well as automatic cleanup, refinement and interpolation features.

When following the different tabs of VolWeaver, the user is guided through a workflow of embedding high-resolution tomograms into low-resolution overview volumes, montaging of adjacent tomograms of equal resolution, stacking of serial tomograms, drawing and clean-up of segmentation masks, semi-automatic measurement of substructure dimensions and per-object volume and surface area (Figure 3B). The program also features a wide range of common filters and data conversion functionalities for tomograms, masks and IMOD-style models.

Embedding, montaging and model to volume alignment use a combination of manual positioning and automatic, intensity-guided fitting of volumes relative to each other. Embedding and montage offer the option to save output as transform files (in JSON format) that can then be loaded into serial join, thus avoiding the build-up of large intermediate files.

Serial joins start with the manual z-cropping of each volume or composite. The user is then guided to the landmark placement where anchor points are defined side-by side to the z-faces of adjacent tomograms and used for the subsequent fitting step that calculates the transformations needed to fit landmarks between volumes. An optional cropping step is recommended when working with very large assemblies, as it can reduce the disk space needed to save the full composite volume.

Modelling in VolWeaver is based on MRC masks but IMOD-style models are supported and can be converted to masks and back for further processing within or outside of VolWeaver. In addition to drawing tools, the VolWeaver modelling tab also contains a range of cleanup options to refine automatic segmentations generated by third-party algorithms, such as deep-learning based segmentation in EMAN2, Dragonfly or MemBrain (*57–59*).

The measurement workflow is the most use-case specific part of VolWeaver that we designed to measure the dimensions of mitochondrial cristae in *P. falciparum*. The workflow is aimed at determining the dimensions of substructures within objects that have been outlined by manual or automatic segmentation. It starts with an optional detection of membrane invaginations, which runs automatically, and can be manually refined by the user (Figure 3C). Based on the derived invagination mask, or alternatively, on manually placed length anchors, length and diameter of the invaginations or any other membrane-enclosed substructure can be determined. The length is directly calculated from the skeleton length path of detected invaginations, or from the anchor derived length path.

From each length path, orthogonal lines can be created at regular increments or a fixed number. They are helper lines that are orthogonal to the length path’s local tangent and align with the z axis. VolWeaver then marches along the orthogonal lines to find the local diameter of the object at that position. This marching approach ensures that the true diameter is found even if the length path is not centred within the invagination. Diameter lines are created orthogonally to the orthogonal line, such that they lie within the xy-plane. We specifically measure in the xy-plane to minimise distortions arising from the missing wedge/cone effect (*60*) that obscures the z-boundaries of objects in TEM-tomography.

In addition to the length and diameter measurements, surface area and inner volume of each object and each invagination are calculated automatically. Surface area is estimated by integrating the gradient magnitude of a Gaussian-smoothed binary mask, an approach related to implicit-surface morphometry and differential-geometric surface estimation methods (*61*). Accuracy of the outputs for length, diameter, volume, and surface area has been validated using synthetic mask variations of a mitochondrion-like object of calculatable geometric dimensions (shape overview and validation results in Supplementary Information 2, summary in Table 1). Error margins of the VolWeaver outputs for object surface area, object volume, invagination length and invagination diameter consistently remained below 5%, irrespective of orientation, anisotropy, gapped segmentation and open ends in z.

**Table 1:** Overview of the validation results for VolWeaver measurements. Error margins remain below 5% compared to the calculated geometric ground truth, irrespective of orientation, anisotropy, gapped segmentation and open z-ends. surface area and volume of the object (SA_obj_ and V_obj_) as well as length and diameter (l_inv_ and d_inv_) of 5 invaginations per validation object have been calculated and measured.

| validation object | relative error (%) |  |  |  |
| --- | --- | --- | --- | --- |
| | $SA_{obj}$ | $V_{obj}$ | mean $l_{inv}$ | mean $d_{inv}$ |
| synthetic mito-like mask (anisotropic, $z = 1.5$) | -0.95 | 0.04 | 0.61 | -1.90 |
| mito-like mask ($z = 1.5$), x-rotated | -0.80 | 0.20 | 4.19 | -1.90 |
| mito-like mask ($z = 1.5$), xy-rotated | 1.18 | 0.29 | 2.60 | -1.66 |
| mito-like mask ($z = 1.5$), xy-rotated, small gaps | 1.15 | 0.29 | 2.38 | -1.66 |
| mito-like mask ($z = 1.5$), xy-rotated, small and large gaps | 1.02 | NaN (leaky) | 3.72 | -1.66 |
| mito-like mask ($z = 1.5$), xy-rotated, small and large gaps, open z ends | -1.91 | NaN (leaky) | 4.47555 | -1.66 |
| mito-like mask (isotropic, $z = 1.0$), xy-rotated | -3.45 | 0.30 | -1.72985 | -2.60 |
| mito-like mask (highly anisotropic, $z = 2.0$), xy-rotated | 4.30 | 0.29 | 4.74645 | 0.92 |
| mito-like mask ($z = 1.5$), y-rotated | 0.68 | 0.27 | 2.3684 | -0.73 |

## CONCLUSION AND OUTLOOK

Together with a carefully optimised protocol for *P. falciparum* sample preparation, we are able to generate large high-resolution datasets with corresponding cellular context and informative quantitative analyses using VolWeaver. The combination of strategic acquisition of low-magnification overview tomograms with high magnification zoom-ins allows for a wide context without sacrificing resolution in relevant areas.

Even though staining and resin embedding reduces the achievable resolution, the opportunity for multiple acquisitions at overlapping ROIs and the option to recover full cell volumes reveals information that, depending on the research question, may be more conclusive than local resolution maximisation.

Even though with VolWeaver we do not offer advanced deep learning-based segmentations ourselves, it can be a useful complementation to such tools, especially where the results of automatic segmentations are promising, yet not accurate enough to use as final output.

VolWeaver also includes modelling and measurement tools, which are optimised for complex membrane architectures such as the ones we observe in *P. falciparum* mitochondria. Measurement tools are aimed at our own use case, but the concept is valid for any other segmentation mask in which length, diameter, volume or surface area of substructures or membrane folds need to be measured. The process is human supervised, which ensures that the user can directly see whether the algorithm works well on their data.

VolWeaver is not only a useful resource for other researchers but it also demonstrates that a new era of software development has begun. Using Claude AI, we were able to vibe code the program within months and thereby sped up our research output, which reduced complexity and compatibility bottlenecks in our data processing workflow.

Our advancements in sample processing may unlock structural studies for other research teams in the parasitology field and are a promising starting point for the processing of other unicellular organisms.

Together, our workflow integrates high-resolution structural information within cellular context, enabling the study of complex biological architectures across scales without sacrificing either detail or completeness.

## MATERIALS AND METHODS

### *P. falciparum* maintenance and sexual conversion

*P. falciparum* asexual blood-stage parasites were cultured in complete RPMI medium (with 25mM HEPES, 100mM hypoxanthine, and 24mM sodium bicarbonate) supplemented with 10% A+ human serum and 5% 0+ human red blood cells (Sanquin, The Netherlands). Parasite cultures were kept in a low oxygen environment (3% O_2_, 4% CO_2_) at 37°C. Culture parasitaemia, was monitored with Giemsa-stained blood smears using a 100x oil objective.

Sexual conversion was induced in a trophozoite-stage culture, adding Albumax-containing medium for 36h and changed every 12h (59). Gametocytes were then cultured with normal culture medium until maturation. During the entire procedure gametocytes were kept at 37°C in low oxygen conditions. To remove ABS from culture, gametocytes were cultured with heparin (20U/ml) for 4 or 5 days.

### Electron microscopy sample preparation

Mature gametocytes were isolated using MACS LS or LD columns. Shortly, mature gametocyte cultures were flushed in the LD columns or in the LS columns attached to a magnet using a 23G needle to regulate the flow. To elute gametocytes, columns were removed from the magnet and washed with incomplete medium. To prevent gametocytes activation during purification, MACS column, magnet and incomplete medium were pre-warmed at 37°C and kept in temperature for the entire procedure. For chemical prefixation, purified gametocytes were incubated in 1% PFA and 0.0075% GA in MOPS buffered saline (10mM MOPS free acid, 2.5mM NaOAc, 137mM NaCl, pH 7.4) for a minimum of 30min and pelleted at 600 x g for 10min. Pellets were vitrified using the Leica EM HPM100 or Leica EMPACT high-pressure freezer in 0.1 x 0.2mm membrane carriers. Water removal and staining was achieved by freeze-substitution in 1% OsO_4_, 0.2% uranyl acetate and 1% water in spectroscopy-grade acetone. Stepwise freeze-substitution consisted of a 48h incubation at -90°C, sloped increase to -60°C within 12h, 10h incubation at - 60°C, sloped increase to -60°C within 12h, incubation at - 30°C for 8 h and removal of UA by washing in 1% OsO_4_ in acetone at -30°C. The samples were then transferred to ice, OsO_4_ was replaced with pure acetone, and the samples were further stained with 1% TA in acetone for 1h and with 1% OsO_4_ in acetone for 1h. Then the samples were embedded in Epon (46% HY 964 Araldite, 39.6% Epoxy Embedding Medium, 13% MNA, 1.4% DMP-30). The Epon-to-acetone ratio was gradually increased (1:2, 1:1, 2:1, 3:1), followed by embedding in pure Epon. Fully infiltrated samples were hardened at 70°C for 5d. Thin sections of 65nm or serial semi-thin sections of 250nm were collected on meshed or single-slot copper grids with formvar/carbon coating. Thin sections were post stained with 2% uranyl acetate in water for 5min and Reynolds lead citrate (Reynolds, 1963) for 2min. Semi-thin sections were labelled with 15nm gold fiducials for tilt-series alignment in IMOD (*53*).

### Special precautions for high-pressure freezing of living cultures

During and after isolation on MACS LS columns (see previous section) gametocytes were kept strictly at 37°C to prevent stress and activation. Sample pelleting was delayed until minutes before freezing and all equipment used during sample placement and freezing was kept on heating plates at 37°C to avoid temperature drops during preparation. Once enclosed in the membrane carrier assembly, samples were high-pressure frozen immediately, using the Leica EMPACT high-pressure freezer in adherence to safety protocols for processing infectious materials (*44*).

### Transmission electron microscopy

Thin sections of 65nm were imaged at the JEOL1400 operating at 120kV. For high-resolution tomographic acquisition, semi-thin sections of 250nm were imaged at the JEOL2100 operating at 200kV, using SerialEM for automated tilt-series acquisition (*54*). Tilt-range was usually kept at +/- 60° with an increment of 1.5° and dual-axis acquisition. For low-resolution overview tomograms we mainly used the JEOL1400 operating at 120kV, with tilt-range of 45° and increments of 2° (dual-axis). Defocus was calculated based on field of view: 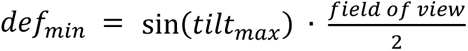

### Data processing

Single-axis micrographs were contrasted in FIJI (*62*). Tilt-series were reconstructed in IMOD (*53*) using batch reconstruction mode with fiducial alignment and dual axis combination. SIRT-like filters at 20 iterations were chosen for most of the high-resolution tomograms, while low-resolution overview tomograms were reconstructed as pure back projections. All processing downstream of reconstruction was performed in VolWeaver.

### General code built of VolWeaver

The entire application was vibe coded in python, using the Claude AI desktop application. The initial framework of VolWeaver was based on a detailed description of the purpose and proposed structure of the user interface and functionality and implemented by Claude Opus 4.6. It included tabs 1 – 3 of the application and had rudimentary functionality at first version. Bug removal, upgrades and the integration of additional modules was performed in an iterative, human supervised process, with the assistance of Claude Sonnet 4.6, and later Opus 4.8 and Sonnet 5. Close supervision and rigorous testing of each function implemented into the code ensured proper functionality and reliability of the software.

### Automatic fitting of embed volumes and montages in VolWeaver

The embed and montage workflows both contain automatic fitting in three modes (translation-only; rigid: translation plus in-plane rotation; and affine: translation, rotation, and anisotropic scaling). In both contexts, one volume serves as the fixed reference, while all other volumes loaded into the canvas are treated as moving volumes whose transforms are optimised independently. Each moving volume is described by a 4×4 affine transformation matrix that maps coordinates in the reference (fixed) volume to corresponding coordinates in the moving volume. The matrix is parameterised by nine degrees of freedom: three translations, three Euler rotation angles, and three anisotropic scale factors. Intensity normalised volumes are bandpass-filtered using a Difference-of-Gaussians kernel with standard deviations σ₁ = 1.0 and σ₂ = 4.0 voxels. The filter suppresses low-frequency intensity gradients while enhancing structural boundaries and membrane densities. An initial translational offset is estimated through amplitude-normalised phase correlation.

For rigid and affine modes, the translation-refined transform is subsequently refined by an exhaustive grid search over in-plane (z-axis) rotation. A total of 41 candidate rotation increments is sampled uniformly across a user-defined search window. For each candidate rotation, the moving volume is re-rendered under the trial transform and the normalised cross-correlation (*NCC*) coefficient is computed. The affine mode extends the rigid solution with a two-stage anisotropic scale search and a final rotation re-check. The total number of NCC evaluations is approximately 250 for affine mode, compared with 41 for rigid and 0 for translation-only mode.

### Surface area measurement in VolWeaver

Object surface area is estimated by the co-area (gradient-integral) method, which is independent of membrane orientation. The binary mask of the object is convolved with a Gaussian kernel that is isotropic in physical space (lateral σ = 0.65 voxels; axial σ = 0.65/ζ voxels, with ζ being the z-scale), and the surface area is obtained as the volume integral of the gradient magnitude of the smoothed mask, SA = ∫|∇Mσ| dV, which equals the area of its 0.5 iso-surface. Gradients are computed by central differences with the axial component scaled by ζ, and the integral is converted to physical units (×ζ, ×p², p being the lateral voxel size).

### Enclosed volume measurement in VolWeaver

The lumen volume enclosed by an object is computed slice by slice. On each axial plane the membrane contour is sealed by a one-voxel dilation (to bridge single-pixel gaps) and its interior filled; the membrane voxels are then excluded so that only interior space is counted, and the summed voxel count is scaled by p³ζ. Leaking fills (through gaps in the membrane or an open contour) are detected per slice by comparing the enclosed area with the area of the membrane’s convex hull: a slice enclosing less than 60% of its hull is flagged as leaking. If more than 5% of an object’s membrane-bearing slices leak, the enclosed volume is reported as undefined (leaky); below that threshold, isolated leaking slices are replaced by linear interpolation between their nearest intact neighbours.

### Invagination detection in VolWeaver

Membrane invaginations are detected on each z-slice as either closed inner lumina or open bays (inward concavities of the sac boundary, recovered as the difference between a morphological closing of the sac envelope and the envelope itself). Per-slice footprints are assembled into three-dimensional invaginations by cross-slice overlap (optionally supplemented by centroid-proximity matching). Each invagination’s volume is derived from the cavity’s voxel count scaled pixel size and z-scaling. The surface area is obtained as described above.

### Automatic length measurement in VolWeaver

A centreline length path is generated from the mouth to the deepest point of each invagination. For path construction, each invagination is reduced to a one-voxel-wide medial axis by three-dimensional skeletonisation and the longest geodesic between medial-axis end points is found with Dijkstra’s algorithm on the skeleton graph using anisotropic edge weights, √(Δx² + Δy² + (ζΔz)²). The mouth end is anchored at the opening in the surrounding membrane and the tip end is extended along its local tangent onto the invagination wall. The length measurement is the anisotropy-weighted arc length of the resulting polyline.

### Diameter measurement in VolWeaver

Diameter measurements are constructed from three automated steps that are built based on the length path. 1) Measurement positions are sampled at uniform intervals along the length path. 2) The local diameter directionality is defined as the xy-aligned orthogonal to the local path tangent. To account for off-centre length paths from manual or inaccurate placement, several measurements are conducted along a z-aligned orthogonal line. To avoid unnecessary measurement of far-off values, measurement z-depth is user-defined and recommended to be chosen equal to the expected object diameter. z-slices are processed in order of increasing distance from the slice closest to the sampling position on the length path. At each sampled z, a diameter chord is grown simultaneously in both the positive and negative diameter directions, until the object wall is reached. A measurement is accepted only when the line crosses the path and contacts an object wall on both sides. Additionally, retainment or rejection of a measurement depends on the combination of the first accepted chord at that position and the user-defined maximum length ratio. The first accepted chord is the first non-zero chord at that position and it is designated the reference sample for that position. Subsequent chord candidates are rejected if their length exceeds the product of the threshold factor and the reference chord length. The reported diameter at each position is the length of the longest accepted chord across all processed z-slices.

### Validation of VolWeaver measurements

To validate length, diameter, volume and surface area measurements, we created a synthetic mask that resembles a typical mitochondrial inner membrane as we see it in *P. falciparum* gametocytes. The object consists of a spherocylinder with five radial blind-ended cylindrical invaginations (main tube: R = 100 nm, L = 600 nm; invaginations: r = 12 nm, depth d = 40 nm). Derivates of the structure included multiple rotations, a range of different z-scales, removal of the caps at both ends of the object’s z-range, and the introduction of small or larger segmentation gaps. Length, diameter, volume and surface area of the object and invaginations were measured with the automatic invagination detection, length path generation and diameter determination workflow in VolWeaver and independently calculated based on the geometrical characteristics of the synthetic mask (results in Supplementary Information 2, summary in Table 1).

### z-gap interpolation of incomplete masks in VolWeaver

Manual segmentations at increments of 2-10 z-slices (depending on the complexity of the object) are interpolated with signed-distance transforms (SDTs). Interpolation is performed independently for each object, while overwriting of existing non-zero values is blocked. Default mode performs SDT directly on the segmentation mask. Membrane mode and invagination sensitive membrane mode refine the process for the respective purpose.

In membrane mode, membranes are recovered as the boundary bands of their flood-filled interior (note that contours must be closed for the flood-fill to be accurate). In invagination sensitive membrane interpolation, the outer sac and the invaginations are reconstructed separately and then combined, so each membrane line has a single origin. The main sac SDT is computed from a smooth envelope in which all invaginations (holes and bays) are filled; the interpolated sac boundary therefore does not itself dip into invaginations. Invaginations are detected as enclosed inner rings or bays in which the main membrane sac dips inward, while the smoothed surface of the deducted solid does not. Both states are detected per slice, as lumen footprints. Footprints are matched by centroid position or xy-overlap to determine whether they are part of the same invagination. The algorithm uses user-defined thresholds for bay and hole size to distinguish between minor membrane irregularities, segmentation errors and true invagination paths. A z-gap cannot be interpolated meaningfully when the invagination configuration changes too greatly between the two drawn slices. Therefore, each interval is scored by a reliability measure equal to the fraction of lumen footprints (over both reference slices) belonging to a bilaterally matched track. Interpolations below user-defined threshold are skipped and flagged for manual inspection.

### Membrane gap bridging of incomplete masks in VolWeaver

All processing is performed slice-by-slice in a user-selected projection plane (xy, xz, or yz) or iteratively across all three planes. While the distance-only mode connects structural endpoints by the shortest straight line, the intensity-guided workflow is based on a filtered volume on which low-cost bridges along high-intensity pixels are identified. In either mode, membrane masks are first skeletonised and line endpoints are defined as path pixels that have exactly one neighbour in the 8-connectivity sense. For intensity-guided gap bridging, two filters are available for membrane enhancement prior to path detection: Hessian of Gaussian and Frangi vesselness (*63*). Each filter option can be finetuned and previewed by the user. For every pair of components for which the nearest anchor pixels are separated by no more than the user-defined maximum gap, a Dijkstra shortest-path search is executed on the filtered image. Bridges are accepted only if their mean per-pixel cost falls below a user-defined threshold sensitivity. The logic of the algorithm retains one last gap, which the user can choose to be closed by Dijkstra search and sensitivity threshold (intensity-guided) or straight Bresenham (distance-only).

## ACKNOWLEDEGMENTS

Our work is supported by funding provided by the Dutch Research Council (NWO; Nederlandse Organisatie voor Wetenschappelijk Onderzoek) (grant numbers OCENW.M.21.087, awarded to T.W.A.K. and L.v.N., and OCENW.XS25.2.029, awarded to I.B.). We thank Stefan Fehr for valuable advice and in AI-assisted coding methods.

## Notes

https://zenodo.org/records/21931154

https://github.com/irinabregy/VolWeaver

### Competing Interest Statement

The authors have declared no competing interest.

