## Supplementary material for "Streamlining large-scale high-resolution electron tomography with VolWeaver": Figure S 1

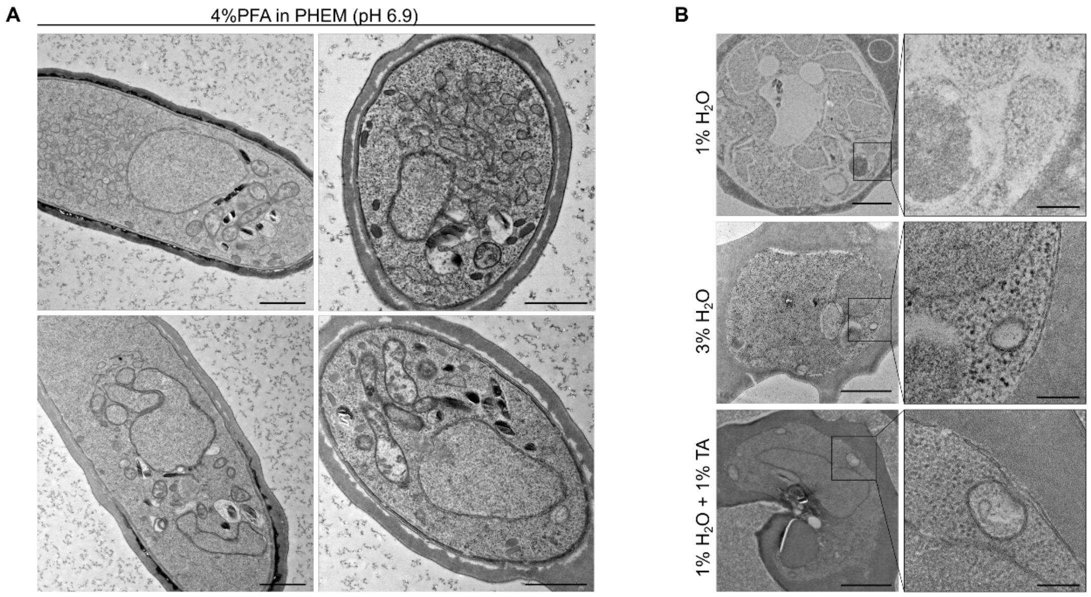


***Figure S 1: Overview of a range of sample treatment options tested during the optimization of* P. falciparum *preparation for TEM.*** *A) Thin section TEM of* P. falciparum *gametocytes prefixed with 4% PFA in PHEM buffer and treated further as described in Figure 1 B. Scale bar: 1µm. B) Thin section TEM of* P. falciparum *blood stage parasites freeze substituted in the presence of 1% or 3% water and (bottom panel) treated with TA after freeze substitution. Scale bars: 1µm / 200nm.*
